# CryoEM structures of MDA5-dsRNA filaments at different stages of ATP hydrolysis

**DOI:** 10.1101/376319

**Authors:** Qin Yu, Kun Qu, Yorgo Modis

## Abstract

Double-stranded RNA (dsRNA) is a potent proinflammatory signature of viral infection. Long cytosolic dsRNA is recognized by MDA5. The cooperative assembly of MDA5 into helical filaments on dsRNA nucleates the assembly of a multiprotein type-I-interferon signaling platform. Here, we determined cryoEM structures of MDA5-dsRNA filaments with different helical twists and bound nucleotide analogs, at resolutions sufficient to build and refine atomic models. The structures identify the filament forming interfaces, which encode the dsRNA binding cooperativity and length specificity of MDA5. The predominantly hydrophobic interface contacts confer flexibility, reflected in the variable helical twist within filaments. Mutation of filament-forming residues can result in loss or gain of signaling activity. Each MDA5 molecule spans 14 or 15 RNA base pairs, depending on the twist. Variations in twist also correlate with variations in the occupancy and type of nucleotide in the active site, providing insights on how ATP hydrolysis contributes to MDA5-dsRNA recognition.

**eTOC:** Structures of MDA5 bound to double-stranded RNA reveal a flexible, predominantly hydrophobic filament forming interface. The filaments have variable helical twist. Structures determined with ATP and transition state analogs show how the ATPase cycle is coupled to changes in helical twist, the mode of RNA binding and the length of the RNA footprint of MDA5.

**Highlights:**

- CryoEM structures of MDA5-dsRNA filaments determined for three catalytic states
- Filament forming interfaces are flexible and predominantly hydrophobic
- Mutation of filament-forming residues can cause loss or gain of IFN-β signaling
- ATPase cycle is coupled to changes in filament twist and size of the RNA footprint

## Introduction

Recognition of viral nucleic acids by innate immune receptors is one of the most conserved and important mechanisms for sensing viral infection. Many viruses deliver or generate double-stranded RNA (dsRNA) in the cytosol of the host cell. RNA duplexes have the A-form double helix structure distinct from the B-form structure of DNA (Pabit et al., 2016), and cytosolic dsRNA is a potent proinflammatory signal in vertebrates. Uninterrupted RNA duplexes longer than a few hundred base pairs (bp) are recognized in the cytosol by the innate immune receptor MDA5 (melanoma differentiation-associated gene-5) (Kato et al., 2008; Kato et al., 2006). The cooperative assembly of MDA5 into ATP-sensitive filaments on dsRNA induces oligomerization of its tandem N-terminal caspase recruitment domains (CARDs) (Berke and Modis, 2012; Peisley et al., 2011). The MDA5 CARD oligomers nucleate the growth of microfibrils of the CARD from MAVS (mitochondrial antiviral signaling protein) (Hou et al., 2011; Wu et al., 2014). The amyloid-like (or prion-like) properties of MAVS CARD microfibrils initiate the assembly and growth of a multimeric signaling platform on the outer mitochondrial membrane, which includes proteins from the TRAF and TRIM families (TRAF2/3/5/6 and TRIM13/25/31) (Hou et al., 2011). The MAVS signalosome activates both type I interferon and NF-κB-dependent inflammatory responses (Hou et al., 2011; Kato et al., 2008; Kato et al., 2006). The assembly of MDA5 filaments on dsRNA also efficiently displaces viral proteins from the RNA while promoting dsRNA-binding and activation of Protein Kinase R (PKR) (Yao et al., 2015), which leads to inhibition of protein translation and hence virus replication (Chung et al.). This effector activity of MDA5 is ATP-dependent but CARD-independent (Yao et al., 2015).

Recently, it was shown that mRNA containing Alu repeats, endogenous retroelements of viral origin constituting 10% of the human genome, can hybridize into long RNA duplexes that must be deaminated by ADAR1 to avoid recognition by MDA5 (Ahmad et al., 2018). A-to-I deamination by ADAR1 destabilizes the of Alu:Alu duplex sufficiently to prevent MDA5 filament formation. Gain-of-function MDA5 mutations or ADAR1 deficiency can cause PKR-mediated translational shutdown and severe autoimmune disorders, including Aicardi-Goutières syndrome (AGS) and other interferonopathies (Ahmad et al., 2018; Chung et al., 2018; Rodero and Crow, 2016).

Crystal structures of MDA5 bound to dsRNA oligonucleotides show that MDA5 binds dsRNA with a modified DExD/H-box helicase core and a C-terminal domain (CTD) (Uchikawa et al., 2016; Wu et al., 2013). The helicase consists of two RecA-like domains, Hel1 and Hel2, and an insert domain, Hel2i, all of which form contacts with phosphate and ribose moieties within the backbones of both RNA strands. The helicase and CTD, linked by a pair of α-helices referred to as the pincer domain, form a closed ring around the RNA (Uchikawa et al., 2016; Wu et al., 2013). The overall structure is similar to those of two other helicases in the same subfamily, RIG-I and LGP2 (Jiang et al., 2011; Kowalinski et al., 2011; Luo et al., 2011; Uchikawa et al., 2016). However, RIG-I and LGP2 use their CTD to bind dsRNA blunt ends, with 5’-di- or triphosphate caps and unphosphorylated, respectively, and both proteins have a much lower propensity than MDA5 to form filaments (Devarkar et al., 2016; Goubau et al., 2014; Luo et al., 2012). A structure of MDA5 bound to viral dsRNA determined by negative-stain electron microscopy at 22 Å resolution showed that MDA5 forms a polar, single-start helix on dsRNA and suggested that in MDA5 the CTD participates in filament formation rather than dsRNA blunt end recognition as in RIG-I and LGP2, but the resolution was insufficient to identify specific intermolecular interfaces (Berke et al., 2012).

The dsRNA binding cooperativity and length specificity of MDA5—which are critical for its signaling activity—are encoded by the filament-forming interfaces, but these remain unknown. To identify this key missing link to gaining a molecular-level understanding of the polymerization-dependent recognition of dsRNA by MDA5, we determined the structures of the MDA5-dsRNA filament by cryo-electron microscopy (cryoEM) in three different helical twist states and three different nucleotide binding states at local resolutions of up to 3.42 Å, allowing us to build and refine atomic models for the filaments. The structures reveal a predominantly hydrophobic pair of filament forming interfaces with the requisite flexibility to accommodate the mechanical properties of dsRNA (Herrero-Galan et al., 2013). Structures bound to ATP, ground state analog AMPPNP, transition state analog ADP-AlF_4_ and no nucleotide show how the ATPase cycle is coupled to changes in helical twist and the mode of dsRNA binding of MDA5. This work shows how MDA5 recognizes long dsRNA ligands and provides a structural basis for its proposed proofreading activity.

## Results

### MDA5-dsRNA filaments have a variable helical twist

MDA5-dsRNA filaments were assembled by incubating recombinant mouse MDA5 with dsRNA in the presence of either ATP, the nonhydrolyzable ATP (ground state) analog AMPPNP, or the transition state analog ADP-AlF_4_. A concentration range of 1 – 10 mM of nucleotide was selected for a favorable trade-off between filament stabilization and vitreous ice formation, while remaining near the physiological range of cellular ATP concentration. Residues 646–663, in flexible surface loop of Hel2i, were deleted to improve solubility, resulting in a 114 kDa polypeptide chain. This “ΔL2” deletion does not interfere with the ATPase, dsRNA binding or interferon signaling activities of MDA5 (Berke et al., 2012; Wu et al., 2013). Filaments were plunge-frozen in liquid ethane for cryoEM data collection. Power spectra from raw images showed meridional reflections confirming the previously determined helical rise of approximately 44 Å for MDA5 within the filaments (Berke et al., 2012) (**Figure S1**). The high average curvature and variability in helical twist of the filaments presented challenges for helical image reconstruction. A cylinder was used as the initial model for 3D image reconstruction, along with the experimentally determined helical symmetry parameters (see **Methods** and (Berke et al., 2012)). To deal with sample heterogeneity, particles were divided into several classes during 3D image reconstruction. Most of the variability between classes was in the helical twist. There was no evidence for discrete twist states, and the number of classes used was arbitrary. Individual filaments contained segments with different twists, with segments of similar twist forming small clusters, indicating that the variability in twist was a local phenomenon (**Figure 1B**). Some twist values occurred more frequently than others and the twist distribution depended on type and occupancy the nucleotide bound (**Figure 1C**). Helical reconstruction of the filaments formed with 1 mM AMPPNP in RELION (He and Scheres, 2017) produced three maps with an overall resolution better than 4 Å (3.68 – 3.93 Å) (**Table S1** and **Figures S2** and **S3**). The maps had distinct helical twists of 74°, 87° and 91°, respectively, but similar rises (43 – 45 Å). With local resolutions up to 3.42 Å, the maps, referred to henceforth as Twist74, Twist87 and Twist91, were sufficiently detailed for atomic models to be built and refined into each map using the crystal structures of human and chicken MDA5 bound to dsRNA oligonucleotides as starting models (Uchikawa et al., 2016; Wu et al., 2013) (**Figures 1D-E** and **S3**). Reconstructions of filaments formed with 2.5 mM AMPPNP, 10 mM ATP, 2 mM ADP-AlF_4_ and without nucleotide, respectively, were subsequently obtained with resolutions of 3.87 – 4.06 Å. Filaments formed with ATP were frozen a few (7 – 8) minutes after addition of ATP to the sample, to prevent ATP hydrolysis from proceeding to completion. Most of the filaments formed with 10 mM ATP had low helical twist (71° – 81°), and most of the filaments formed without nucleotide had high helical twist (86° – 96°). The filaments formed with ADP-AlF_4_ had a narrower distribution of intermediate twists (81° – 91°) (**Figure 1C**). Filaments formed with 2.5 mM AMPPNP had twists spanning a similarly broad range as with 1 mM AMPPNP (71° – 96°).

**Figure 1.**
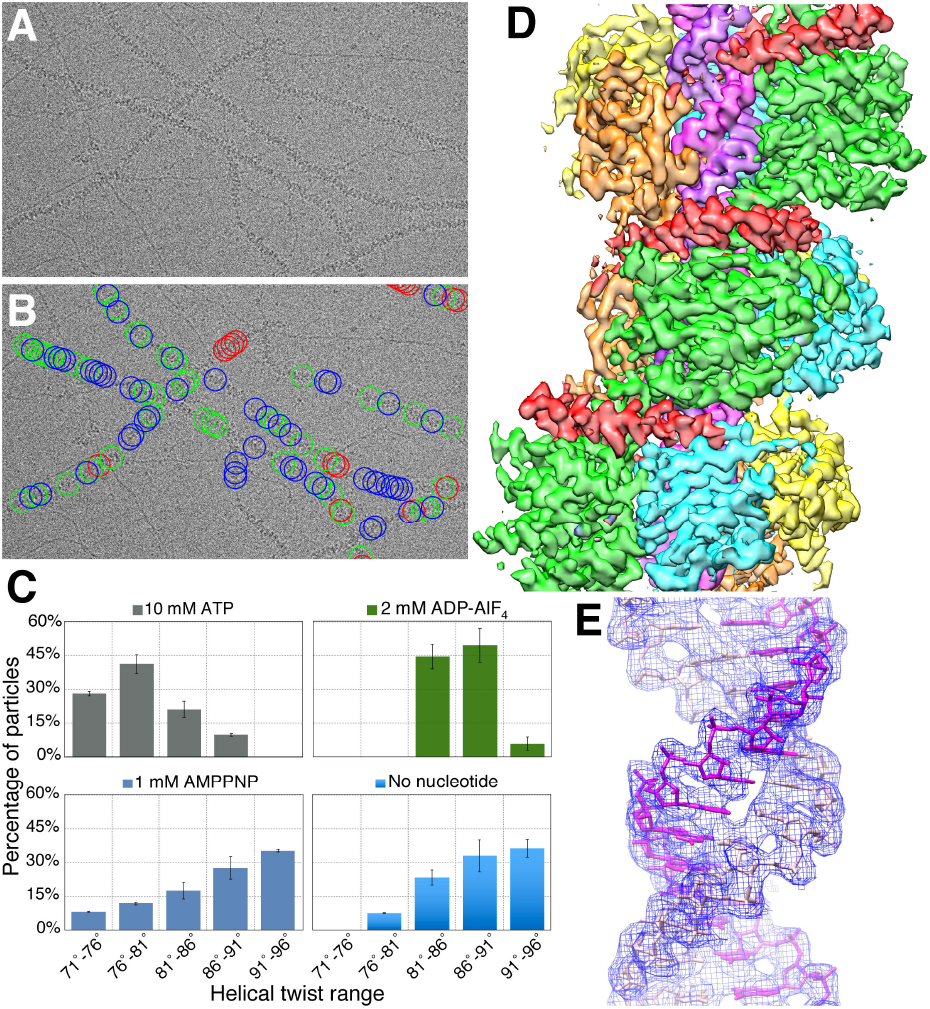
CryoEM image reconstruction of MDA5-dsRNA filaments with helical symmetry averaging. (**A**) Representative electron micrograph of MDA5-dsRNA filaments embedded in vitreous ice. (**B**) Electron micrograph shown in *(A)* with circles drawn around the boxed filament segments that were extracted and used in the helical reconstructions. The circles are color-coded according to the 3D class that they contributed to. Segments that contributed to the structure with a helical twist of 74° (Twist74) are in red. Segments that contributed to the Twist87 and Twist91 structures are in green and blue, respectively. (**C**) Histogram showing the distributions of extracted filament segments as a function of helical twist for the 10-mM ATP, 2-mM ADP-AlF_4_, 1-mM AMPPNP and nucleotide-free datasets. The distributions shown are from 3D classification performed with ten classes per dataset. Error bars represent standard deviation from the mean between replicate 3D classification calculations; n = 3. (**D**) 3D density map of the Twist74 MDA5-dsRNA filament at 3.68 Å overall resolution. The components are colored as follows: Hel1, light green; Hel2, cyan; Hel2i, yellow; pincer domain, red; C-terminal domain (CTD), orange; RNA, magenta. (**E**) The dsRNA density in the Twist74 filament (blue mesh) is shown with the fitted atomic model (in magenta and pink ball-and-stick representation).

### Overall structure of the MDA5-dsRNA filaments

In the refined atomic models presented here, MDA5 forms a closed ring around the RNA (**Figure 2** and **Movie S1**). The low-twist structures (71° – 81°) contain 14 bp of dsRNA in the asymmetric unit contains and density for nucleotide bound in the ATP binding site. The intermediate-twist (81° – 91°) and high-twist (91° – 96°) structures have 15 bp of dsRNA and no interpretable density in the catalytic site. The nucleotide density ranges from absent or very weak in the nucleotide-free structures and the 1-mM AMPPNP low-twist structure (Twist74), respectively, to very strong in the intermediate-twist ADP-AlF_4_ structure. The 2.5-mM AMPPNP and 10-mM ATP structures have nucleotide densities of intermediate strength (**Figure 3**). The nucleotide density in the 10-mM ATP structure is unambiguously more consistent with ATP than with ADP or ADP:Mg^2+^ (**Figure 3C**), suggesting that the low-twist filament segments used in the 10-mM ATP reconstruction predominantly contained ATP that remained unhydrolyzed when the sample was frozen. The Hel1 and Hel2 domains are in the semi-closed state, as defined by (Uchikawa et al., 2016) in all structures except the structure with ADP-AlF_4_ bound, which is in the closed state. Two out of six ATP-binding helicase motifs, motifs Q and I as defined by (Jiang et al., 2011), are engaged with nucleotide in Twist74 and the other low-twist semi-closed structures. All six ATP-binding helicase motifs—motifs Q, I, II, III, Va, and VI—are engaged with ADP-AlF_4_ in the closed structure.

**Figure 2.**
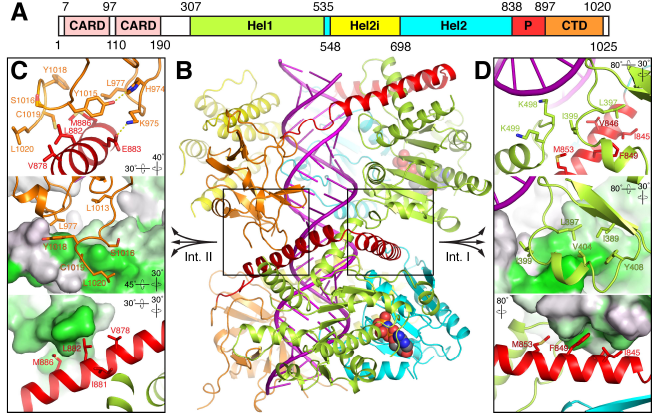
Atomic model of the MDA5-dsRNA filament. (**A**) Domain structure of mouse MDA5. CARD, caspase recruitment domain; Hel1 and Hel2, first and second RecA-like helicase domains; Hel2i, Hel2 insert domain; P, pincer domain; CTD, C-terminal domain. The same color code and domain abbreviations are used in subsequent panels and in Figures 1 and 7A,D. (**B**) Overview of the refined atomic model of the MDA5-dsRNA filament. Two adjacent MDA5 subunits and 28 bp of dsRNA are shown from the structure with 74° helical twist (Twist74). RNA is in magenta. The bound AMPPNP molecules are shown in sphere representation. The two filament forming interfaces are boxed and labelled “Int. I” and “Int. II”. (**C** and **D**) Close-up views showing the key filament forming contacts of Interface II (**C**), and Interface I (**D**). The top panels show the side chains of the residues forming key contacts, with hydrogen bonds shown as yellow dashed lines. In the middle panels the lower protomer in *(B)* is shown in surface representation colored by hydrophobicity from grey to green, with green being the most hydrophobic. In the lower panels, the upper protomer in *(B)* is shown in surface representation colored by hydrophobicity. The orientation of the view relative to *(B)* is indicated for each panel.

**Figure 3.**
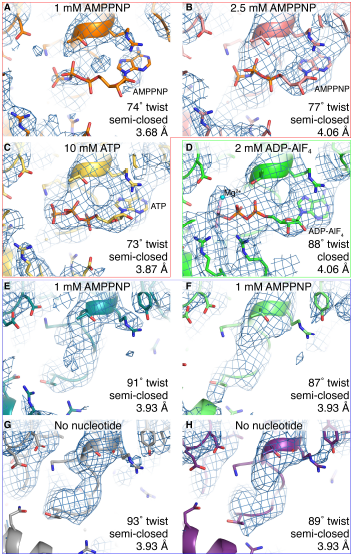
Close-up views of the cryoEM densities and atomic models around the ATP binding site for the selected reconstructions with different helical twists and bound nucleotides. A contour level of 4.5 σ in PyMol (Schrödinger) was used for all eight panels. The AMPPNP, ATP and ADP-AlF_4_ molecules, and selected protein side chains are shown in ball-and-stick representation. The local resolution of the maps is relatively low in this region (see **Figure S3**). Density consistent with a nucleotide triphosphate molecule is visible in the low-twist structures with 2.5 mM AMPPNP (**B**) and 10 mM ATP (**C**), but only weak density is visible in the low twist (74°) 1 mM-AMPPNP structure (**A**) (red box). In the presence of 2 mM ADP-AlF_4_ (**D**) strong density is visible for the nucleotide and aluminium fluoride moiety (green box). The aluminium fluoride is shown in pink and grey and a coordinated magnesium ion in cyan. In the presence of 1 – 2.5 mM AMPPNP or in the absence of added nucleotide (**E**-**H**) there is no interpretable nucleotide density in the catalytic site of the structures with mid- to high helical twist (81 – 96°) (blue box).

The asymmetric units of the three twist classes of filaments have a similar overall structure to the dsRNA-bound crystal structures of MDA5 (**Figure S4**). However, the cryoEM structures contain several novel features, most notably the C-terminal tail of the CTD (residues 1014-1020), which extends towards an adjacent subunit and forms key filament contacts. The cryoEM structures also contain a short acidic loop in Hel1 (residues 428-430), the linker between the pincer domain and the CTD (residues 894-899), and part of an extended loop within the central β-sheet of the CTD (residues 945-949). These features are not present the dsRNA-bound MDA5 crystal structures. Some of these features are present but have different conformations in the crystal and NMR structures of the MDA5 CTD alone (PDB 3GA3; (Takahasi et al., 2009)). The CARDs, which were present in the imaged samples, were not visible in any of the density maps after helical symmetry averaging, even at low contour levels, indicating that the CARDs do not share the helical symmetry of the MDA5-dsRNA filament. Moreover, since MDA5 CARD oligomers could not be distinguished in the raw micrographs, we conclude that CARD oligomers are no larger than approximately 100 kDa, the current size limit for cryoEM single particle imaging, which corresponds to the tandem CARDs of six to eight MDA5 molecules.

### Identification of the filament forming interfaces

The cryoEM structures identify the filament forming surfaces of MDA5. The vast majority of the contacts are hydrophobic and can be grouped into two interfaces, both involving the pincer domain (**Figure 2**). Interface I is formed by a loop in Hel1 (residues 395-408), which forms an extensive set of hydrophobic interactions with the first α-helix of the pincer domain (residues 839-864) and an adjacent loop in Hel1 (residues 497-500) of the adjacent subunit. The loop forms a surface with a shape highly complementary to that of the adjacent pincer helix (**Figure 2D** and **Movie S2**). The most notable interactions are hydrophobic side chain contacts between Leu397, Ile399, Ser400, Glu403 and Val404 in Hel1, and Glu842, Ile845, Val846, Phe849, and Met853 in the pincer helix. These side chains form the core of interface I, with intrinsic structural flexibility arising from the hydrophobic nature of the contacts. The surface area buried by the interface is 452 – 570 Å^2^ in the semi-closed structures and 413 Å^2^ in the closed ADP-AlF_4_-bound structure.

Interface II is formed by the C-terminal tail, which extends from the CTD to form hydrophobic contacts with the second pincer helix (residues 866-891) of the adjacent subunit (**Figure 2C** and **Movie S3**). The core contacts of the interface are hydrophobic side chain contacts between Asp1014, Tyr1015, Tyr1018 and Cys1019 in the C-terminal tail, along with Gly976 and Leu977 in the CTD, and Gln879, Leu882, Glu883 and Met886 in the pincer domain. Glu883 is also positioned to forms hydrophilic contacts with either Lys975 or Gly976, depending on the twist class. The C-terminal tail was not resolved in crystal structures of human and chicken MDA5 bound to dsRNA oligonucleotides, indicating that the structure of the C-terminal tail observed by cryoEM is dependent on head-to-tail intersubunit filament contacts. The surface area buried by the interface II is 413 – 433 Å^2^ in the semi-closed structures and 471 Å^2^ in the ADP-AlF_4_-bound structure. Thus, whereas interface I buries a larger area than interface II in all three twist classes with ATP or no nucleotide bound, the opposite is true in the ADP-AlF_4_ structure, which has intermediate twist.

The residues listed above as forming interfaces I and II are generally either conserved or similar in MDA5 but not RIG-I sequences from vertebrate species (**Figure S5**). The following residues are strictly conserved across terrestrial vertebrates in MDA5 but not RIG-I: 397-403, 497-500, and 846 in Interface I; and 879, 883, 886, 976, 977, 1015 and 1019 in Interface II.

A third minor filament contact point is formed by Met571 in Hel2i, which forms hydrophobic side chain contacts with Glu773 from the Hel2 domain of the adjacent subunit (**Figure 5A**). The contact area of this interface is small, 141 Å^2^ in Twist74, and only 42 – 65 Å^2^ in the higher-twist classes, representing 4 – 13% of the total filament interface area. This interface is absent the ADP-AlF_4_-bound structure, and Met571 is not strictly conserved in mammalian MDA5 sequences.

### Flexible MDA5 interfaces lead to variable helical symmetry

The extent to which the MDA5 filament-forming interfaces are predominantly hydrophobic in nature is striking. Hydrophobic interfaces can be intrinsically structurally flexible and allow intersubunit rotation in the absence of the chemical and geometric restraints imposed by polar hydrogen bonds or salt bridges (Li et al., 2005). Comparison of the filament forming interfaces in the different twist classes shows that although the interfaces are broadly conserved, there are significant differences in how they form in each twist class. Superposition of the Twist74 and Twist91 structures using a the Hel1 domain of a specific subunit as the reference (Rmsd 0.75 Å) highlights how variable and flexible the filament-forming interfaces are, with differences of up to 20 Å in the resulting positions of the domains of adjacent filament subunits (**Figure 4**). This large variability in the orientation of the adjacent subunits within the filament is possible specifically due to the structural versatility of the Hel1 interface loop component of Interface I (residues 395-408) and of the C-terminal tail component of Interface II (residues 1014-1020). These components have significantly different conformations in each twist class, each adapting their structure to maintain hydrophobic contacts with the apposed pincer domain helix from the adjacent subunit (**Figures 4E-F** and **S6**). The interface components within the pincer domain, constrained by helical secondary structure, have the same local conformation in the different twist classes, though the pincer helices, which pack tightly onto the Hel1 and Hel2 domains, follow the larger shifts in domain positions mentioned above. The specific hydrophobic contacts formed in the different twist classes are similar but not identical. For example, the C-terminal tail forms contacts two helical turns further down the second pincer helix in the Twist74 structure than in the Twist91 structure, and the Hel1 interface loop adopts a different conformation in Twist74 than in the other cryoEM and crystal structures (**Movie S4**). The net result is that the Hel1 interface loop functions as a flexible finger and the C-terminal tail as a flexible arm, allowing similar but not identical sets of hydrophobic contacts to be maintained between subunits. This bears similarity to the capsid proteins of some spherical viruses, which have flexible C-terminal arms and internal loops that allow the quasi-equivalent assembly of capsomeres in slightly different symmetry environments within the icosahedral lattices of viral capsids (Liddington et al., 1991).

**Figure 4.**
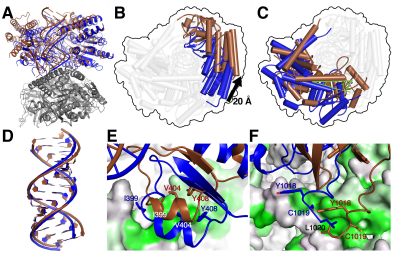
Differences in the relative domain positions and filament contacts in the cryoEM structures with 74° and 91° helical twists. (**A**) Overview of two protomers of the Twist74 and Twist91 structures superimposed on each other using the pincer domain of the lower protomer (grey) as the reference. The upper protomer of Twist74 is in blue and that of Twist91 is in brown. (**B** and **C**) Top views along the helical axis of the upper protomer from panel *(A)* showing the shifts in the positions of Hel2i and CTD (**B**), and Hel1, Hel2 and pincer (**C**). The 20 Å translation in Hel2i and 12° rotation in the pincer domain helices are highlighted. The highlighted domains are colored as in the upper protomer in *(A)* and the remaining domains are shown in transparent grey representation for clarity. The outer contour of the superimposed structures is shown for reference as a black outline. (**D**) Close-up of the dsRNAs from the Twist74 and Twist91 structures from the structural alignment in panel *(A).* The root mean square deviation of the atoms in the 14 superimposed RNA base pairs is 1.23 Å. (**E** and **F**) Close-up views of filament interface I (**E**) and interface II (**F**). The protomer of Twist74 shown in grey in *(A)* and used as the alignment reference is shown in surface representation colored by hydrophobicity as in Figure 2. Key interface residues in the adjacent protomer are shown in ball-and-stick representation with Twist74 in blue and Twist91 in brown.

### Functional importance of MDA5 interfaces in cell signalling

The filament interfaces are critical to MDA5 function because they encode the dsRNA binding cooperativity of MDA5, which promotes binding of MDA5 molecules adjacent to MDA5 molecules already bound to RNA rather than binding to a new site on dsRNA (Berke and Modis, 2012; Peisley et al., 2011). Hence the filament interfaces encode the propensity of MDA5 to form long filaments on RNA and signal more actively from longer dsRNAs. To test the functional importance of the filament forming interfaces revealed in our cryoEM structures, we examined the effect of structure-based mutations targeting the interfaces in cell signalling assays and filament forming assays. To assay cell signaling activity, a panel of sixteen expression constructs encoding human MDA5 variants was generated, with each variant bearing one, two or three structure-based point mutations at one of the filament forming interfaces. The plasmids were individually co-transfected into HEK cells together with plasmids encoding firefly luciferase under control of the IFN-β promoter and *Renilla* luciferase under a constitutive promoter. After expression for 6 h, cells were transfected with polyI:C RNA to induce MDA5 signaling. IFN-β-dependent induction of firefly luciferase was measured in cell lysates 24 h post-induction, as firefly luciferase luminescence normalized against *Renilla* luciferase luminescence (**Figure 5**). The expression level of each MDA5 variant in HEK cells was assessed by Western blotting (**Figure 5C**). With a few exceptions discussed below, the mutations tested significantly reduced or abolished luciferase signaling. The following mutations abolished signaling completely, reducing luciferase activity from the 16-fold induction seen with WT MDA5 down to background levels: T497A/K498A/Q499A and D848A/F849A, targeting key interface I contacts listed above. L396A/K397A/I398A, targeting the Hel1 interface loop, resulted in a 6-fold reduction in luciferase activity (some residue numbers are different in human and mouse MDA5, see **Figure S5**). Deletion of the C-terminal tail (ΔC12) and CTD mutations K975D/D987A, targeting interface II contacts, resulted in a 4-fold reduction in signaling (**Figure 5B**). A triple point mutation in the C-terminal tail (D1014A/Y1015A/E1017K), also targeting interface II, caused a more modest 2- to 3-fold reduction in signaling (**Figure S7B**). The M886A mutation caused a 30% reduction in signaling. The E883R/K884A and K885A mutations, targeting interface II contacts with the C-terminal tail and CTD, had no effect on signaling.

**Figure 5.**
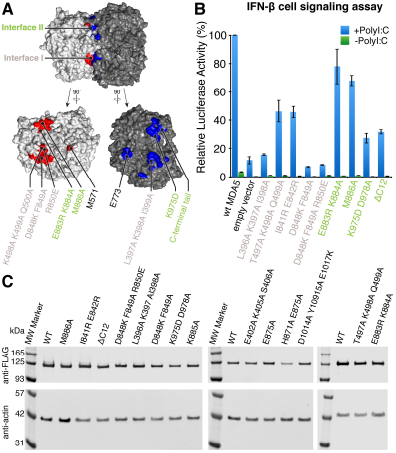
Mutations at the filament forming interfaces abolish or reduce cell signaling in response to dsRNA. (**A**) Location of the filament interface mutations assayed for cell signaling activity in *(B)* and for filament formation in Figure 6. Two filament protomers are shown in surface representation with the filament forming contact surfaces of each protomer colored in red and blue, respectively. The protomers are shown assembled with the helical axis horizontal (top); and separately after being opened like a book with 90° rotations in opposite directions to show the interface surfaces (bottom). Residue labels are colored pink for interface I and green for interface II. Residue numbers refer to the mouse MDA5 sequence. Contact areas were calculated with PISA (Krissinel and Henrick, 2007). (**B**) IFN-β reporter cell signaling assay. Plasmids encoding mutant human MDA5 variants were individually co-transfected into HEK239T cells with plasmids encoding firefly luciferase under an IFN-β-inducible promoter and *Renilla* luciferase under a constitutive promoter. Cells were subsequently transfected with polyI:C RNA (+PolyI:C) or with DMEM (-PolyI:C). Relative luciferase activity was calculated as the ratio of firefly luciferase luminescence to *Renilla* luciferase luminescence. Residue numbers refer to the human MDA5 sequence. Error bars represent standard deviation from the mean between measurements; n = 3. See **Figure S7** for cell signaling data for additional mutants.

The double mutant I841R/E842R, was shown previously to partially inhibit filament formation of human MDA5 on 1-kb dsRNA and to reduce binding affinity for 112-bp but not 15-bp dsRNA (Wu et al., 2013). This pair of mutations maps to the periphery of the pincer helix component of interface I, with Glu842 forming hydrophobic contacts with Thr395 or Leu397 in the Hel1 interface loop. Ile841 does not form any interface contacts, however, and is not conserved in the mouse and chicken sequences. The I841R/E842R mutation resulted in a 2.5-fold reduction of signaling in our assay, consistent with a moderate role of these residues, most likely Glu842, in filament formation (**Figure 5B**). The variant E402A/K405A/S406A had 77% of the signaling activity of WT MDA5, suggesting that the C-terminal portion of the Hel1 interface loop (residues 395-408) plays only a minor role in filament formation (**Figure S7B**).

Two variants unexpectedly caused slight increases in signaling activity: E875A and H871A/E875A (**Figure S7B**). His871 and Glu875 are in the second pincer helix, and their side chains form an intramolecular hydrogen bond in the Twist74 and Twist91 structures. The two residues form interface II contacts only in the Twist87 and Twist91 structures, with the C-terminal tail of the adjacent subunit. Filaments formed by the E875A variant formed aggregates (**Figure S7A**). We speculate that the increase in signaling upon loss of this glutamate side chain, which is conserved in terrestrial vertebrates, may arise from the collapse of filaments into three-dimensional filamentous aggregates.

### Interface mutations that impair signaling hinder polymerization

Cell signaling data show that mutation of MDA5 filament forming residues predominantly results in loss interferon signaling activity, though a few mutations cause slight increases in signaling. To determine whether these changes in signaling activity were due to corresponding changes in the efficiency of filament assembly, we purified a panel of fifteen recombinant mouse MDA5 variants bearing mutations selected from the panel of filament interface mutations assayed for cell signaling in human MDA5. All variants were expressed in similar quantities as WT MDA5 and had similar solubilities and hydrodynamic radii consistent with a monodisperse monomeric population (**Figure S7C-D**). We can therefore exclude the possibility that the loss of signaling activity observed in most interface mutants was due to gross destabilization of the protein fold, which the mutations were designed to avoid. We then assessed the ability of each variant to assemble into filaments on 1 kb dsRNA in the presence of 1 mM AMPPNP, using filament formation in negatively stained electron micrographs as the readout (**Figure 6A**). We found a strong correlation between loss of signaling activity and loss of filament formation. Variants bearing mutations that completely abolished signaling or reduced signaling by at least fourfold (including K498A/K499A/Q500A, D848A/F849A, L397A/K398A/I399A, K975D/D987A and ΔC12) all failed to form filaments. D1014A/Y1015A/E1017A, which caused a 2.5-fold reduction in signaling also failed to form filaments. T841R E842R, which also caused a 2.5-fold reduction in signaling, formed filaments half as long as WT MDA5 (**Figure S7A**). Amongst the least impaired variants, E403A/K406A formed filaments that were 27% shorter that WT MDA5, correlating well with the 23% reduction of signaling observed for the related mutant E402A/K405A/S406A.

**Figure 6.**
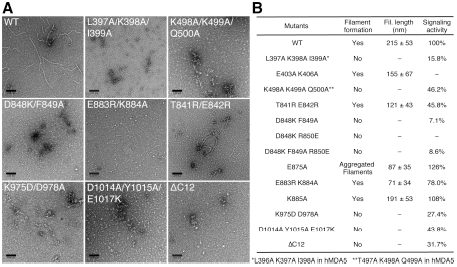
Interface mutations that impair signaling also impair filament formation. (**A**) Representative electron micrographs of WT MDA5 and filament interface mutants in the presence of 1 kb dsRNA, 1 mM AMPPNP and 5 mM MgCl_2_. Scale bars are 100 nm. Residue numbers refer to the mouse MDA5 sequence. (**B**) Table summarizing the filament formation activity, filament length and cell signaling activity of WT MDA5 and selected mutants. See **Figure S7** for electron micrographs of mutants not shown in *(A)*. Residue numbers refer to the mouse MDA5 sequence. For variants that have different residue numbers in the human sequence, the corresponding mutation is listed with human residue numbers at the bottom.

To determine whether filament formation was strictly dependent on dsRNA, we imaged WT MDA5 and several of the mutants in the absence of dsRNA and with 1 mM AMPPNP. None of them formed filaments, including the hyperactive variant E875A, which remained monomeric in the absence of dsRNA (**Figure S7C**). In conclusion, mutation of MDA5 filament forming residues predominantly results in loss of cellular MDA5 signaling activity. Most mutations inhibit MDA5 filament formation and abolish cellular interferon signaling activity, but one pair of mutations appears to increase signaling activity by promoting the collapse of filaments into filamentous aggregates.

### MDA5-RNA contacts and how they vary across the twist and nucleotide states

The dsRNA binding interfaces vary in the different twist classes. The protein-RNA contact area decreases as twist increases, with 2,315 Å^2^ for Twist74, 2,102 Å^2^ for Twist87 and 1,926 Å^2^ for Twist91. The ADP-AlF_4_-bound structure, which has a twist of 88°, has a protein-RNA interface of 2,152 Å^2^, similar to Twist87. The number of contacts decreases with increasing twist accordingly. The CTD forms stronger RNA contacts in the Twist74 structure than in the Twist91 structure (849 Å^2^ versus 471 Å^2^ contact area), whereas the opposite is true for the Hel2i domain (293 Å^2^ versus 321 Å^2^ contact areas). The intermediate-twist ADP-AlF_4_-bound structure is more similar to Twist74 in how its CTD and Hel2i domain bind RNA, with contact areas of 751 Å^2^ and 231 Å^2^, respectively. Within Hel2i, Gln581 is positioned in the Twist87 and Twist91 structures to form hydrogen bonds with either or both bases of an RNA base pair. Within the CTD, Ile923, Glu924 and Met926 form hydrophobic contacts with the RNA backbone in the Twist74 structure; the side chain of His927 forms hydrogen bonds with the O2 or N2 atom of a pyrimidine base (U or C, respectively), and with the ribose hydroxyl group (the latter is present in all three twist classes). The CTD capping loop (residues 944-953), so-called because it binds to RNA blunt ends in related helicases RIG-I and LGP2 (Li et al., 2009; Pippig et al., 2009; Wang et al., 2010), is disordered in the Twist87 and Twist91 structures and in the MDA5-dsRNA crystal structures but partially ordered in the Twist74 structure, in which it forms contacts with the dsRNA backbone at the minor groove as had been predicted (Uchikawa et al.). By contrast, in the crystal and NMR structures of the MDA5 CTD only (PDB 3GA3; (Takahasi et al., 2009)) the capping loop and flanking residues 947-953 extend the central β-sheet of the CTD, forming an additional pair of β-strands. The resulting conformation is incompatible with dsRNA binding, suggesting that dsRNA binding causes the capping loop to peel off from the central β-sheet.

There is less variation across the cryoEM structures in the number of RNA contacts formed by the helicase motifs. Eight out of the ten RNA-binding helicase motifs are engaged with the RNA (motifs Ia, Ib, Ic, IIa, IV, IVa, V and Vc, as defined by (Jiang et al., 2011), with IVb and Vb not engaged). Hel2 forms a more extensive set of contacts than Hel1, with contact areas of 565-660 Å^2^ for Hel2 versus 492-502 Å^2^ for Hel1. Many of the Hel2-RNA contacts are formed by the Hel2 loop (residues 758-767, adjacent to RNA-binding helicase motif IVa), identified previously as a key element for dsRNA stem recognition by inserting into the major groove (Wu et al., 2013). This loop has a similar conformation in the cryoEM and crystal structures (Uchikawa et al., 2016; Wu et al., 2013), and insertion of the loop into the RNA major groove causes a similar widening of the groove from 12 Å to 16 Å. The groove is widened further, to 18 Å, in the ADP-AlF_4_-bound structure. Moreover, in the Twist74 and ADP-AlF_4_-bound structures, His759, at the apex of the Hel2 loop, is positioned so that it could form a hydrogen bond with the O4 or N4 atom of a pyrimidine base. The analogous residue in chicken MDA5 (His733) forms a hydrogen bond with an RNA base, albeit with a purine (G), in a crystal structure in complex with ADP:Mg^2+^ (Uchikawa et al., 2016). In contrast, in the Twist87 and Twist91 structures the Hel2 loop does not form any base contacts and is less firmly engaged with the RNA major groove. Previous work has shown that the Hel2 loop is required for dsRNA-dependent ATP hydrolysis by MDA5 (Wu et al., 2013). Hence, our data support the hypothesis that ATP binding and progression of catalysis to the transition state promote progressive insertion of the Hel2 loop into the RNA major groove, causing a widening of the groove (**Movie S5**). Consistent with this, the dsRNA is slightly stretched along its helical axis in the cryoEM structures relative to free dsRNA (Pabit et al., 2016), and the increase in the rise per bp over the asymmetric unit versus free dsRNA is 13% in the ADP-AlF_4_-bound structure versus only 8% in the Twist87 structure, which has very low nucleotide occupancy in the active site.

### ATP hydrolysis causes Hel1-Hel2 rotation, increasing RNA footprint and filament twist

The structure of MDA5-dsRNA filaments bound to the catalytic transition state analog ADP-AlF_4_ captures a key intermediate in the ATPase cycle. The Hel1 and Hel2 domains are in the fully closed state, with all six nucleotide binding motifs correctly positioned for catalysis. The configuration of the helicase and pincer domains is similar to that in the closed structures of RIG-I and LGP2 in complex with ADP-AlF_4_ (Kowalinski et al., 2011; Uchikawa et al., 2016). Superposition of the ATP- and ADP-AlF_4_-bound MDA5 structures via Hel1 shows that, as in RIG-I and LGP2, the transition from the semi-closed state to the closed state involves a 6 – 7° rotation of Hel2 relative to Hel1, which brings nucleotide binding motifs Va and VI in Hel2 into position for catalysis (**Figure 7A**, **Movie S5**). This rotation of Hel2 relative to Hel1 is transduced by the pincer domain and is part of the conserved allosteric mechanism coupling ATP hydrolysis to dsRNA binding in RLRs (Rawling et al., 2014). As in LGP2 (Uchikawa et al., 2016), the transition from the semi-closed ATP-bound state to the closed ADP-AlF_4_-bound state is accompanied by a shift in the interactions with the RNA backbone made by Hel2 (via motifs IVa and V) by one phosphate along the backbone in the 3’ direction, whereas the interactions of Hel1 with the RNA (via motifs Ia, Ib, Ic, and IIa) are unchanged. The net effect of this shift in Hel2-RNA backbone interactions is a 1-bp increase in the overall footprint of MDA5, from 14 bp in the ATP-bound state to 15 bp in the transition state. Hence, progression from the catalytic ground state (semi-closed, ATP-bound) to the transition state (closed, ADP-AlF_4_-bound) induces a rotation of Hel2 relative to Hel1 that increases the RNA binding footprint by 1 bp. We conclude that ATP hydrolysis by MDA5 is directly coupled to a 1-bp increase in RNA binding footprint.

**Figure 7.**
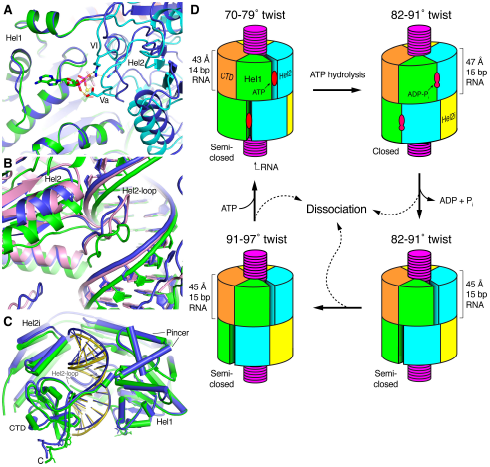
Comparison of the closed ADP-AlF_4_-bound structure with the semi-open structures and schematic model of the ATPase cycle and proofreading mechanism of MDA5. (**A**) Superposition of the Twist74 AMPPNP-bound structure (blue) on the ADP-AlF_4_-bound structure (colored by domain as in Figure 2) using the Hel1 domain as the reference. A close-up view of the nucleotide binding site and Hel1-Hel2 domain interface is shown. The nucleotide binding motifs Va and VI are labeled. Only the ADP-AlF_4_ nucleotide is shown, for clarity (in ball-and-stick representation). (**B**) Superposition of the Twist74 (blue) and Twist87 (pink) AMPPNP-bound structures onto the ADP-AlF_4_-bound structure (green) using the Hel1 domain as the reference. A close-up view of the Hel2-loop and its interactions with the dsRNA is shown. See also **Movie S5**. (**C**) Superposition of the Twist74 AMPPNP-bound structure (blue) on the ADP-AlF_4_-bound structure (green) using the Hel1 domain as the reference. The RNA is shown in darker shades of blue and green. (**D**) Model of the ATPase cycle and proofreading mechanism. Only two filament protomers are shown for clarity. The low-twist (71 – 81°) structures correspond to the ATP-bound catalytic ground state, the intermediate-twist (81 – 91°) ADP-AlF_4_-bound structure is the transition state, and the intermediate- (81 – 91°) and high-twist (91 – 96°) states represent nucleotide-free states. The four panels related to the panels in **Figure 3C-F**.

Along with the increase in RNA binding footprint, the rotation of Hel2 relative to Hel1 upon ATP hydrolysis causes a shift in the position of the RNA backbone at the Hel2 contact site in the direction of the Hel2 rotation. As noted above, the Hel2 loop forms many of the Hel2-RNA contacts, inserts into the RNA major groove and causes widening of the groove. The extent of the widening is greater in the transition state, which contributes to the shift in RNA backbone position relative to the ground state. The net observed shift of the RNA backbone in the transition state suggests that ATP hydrolysis causes a local distortion of the RNA at the Hel2 loop contact site (**Figure 7B**, **Movies S1-S5**). Consistent with this, in the mid- and high-twist semi-closed structures (Twist87 and Twist91) the Hel2 loop is less firmly engaged with the RNA major groove, the RNA backbone is in a similar position as in the ground state (**Figure 7B**), and the overall helical rise is reduced by 2 Å relative to the transition state (**Table S1**). Hence the RNA is less distorted in the low nucleotide-occupancy Twist87 structure than in the transition state, which has the same twist. This suggests that the strain introduced in the RNA in the transition state dissipates upon dissociation of the nucleotide and relaxation of the Hel1-Hel2 interface back to the semi-closed state. Notably, the Hel2-RNA backbone interactions of the Twist87 and Twist91 structures are in the same register as in the ground state structure (Twist74) despite having the same 15-bp footprint as the transition state structure. This suggests that the increases in RNA binding footprint and helical twist gained upon ATP hydrolysis are maintained after dissociation of the nucleotide.

In contrast to LGP2 and RIG-I, the conformational change from the ATP-bound ground state to the ADP-AlF_4_-bound transition state is coupled to shifts in both the Hel2i and CTD domains such that the transition state structure wraps more tightly around the dsRNA, reducing the gap between Hel1 and the CTD by approximately 5 Å (**Figure 7C**). This slightly tighter winding of MDA5 around the RNA could generate overwinding of the RNA, which could explain the increase in helical twist that occurs in going from the ground state to the transition state.

## Discussion

The dsRNA binding cooperativity and length specificity of MDA5, which are critical for its signaling activity, are encoded by the filament-forming interfaces. The cryoEM structures of MDA5-dsRNA filaments determined in this study identify two filament forming interfaces, providing an essential missing link to understanding the polymerization-dependent recognition of dsRNA by MDA5 at the molecular-level. The predominantly hydrophobic nature of the filament-forming contacts provides the flexibility necessary to support cooperative filament assembly on inherently flexible dsRNA duplexes. We have shown that mutation of filament forming residues results in loss of filament formation and MDA5-dependent signaling, with the exception of a pair of mutations, which moderately enhances signaling. In a clinical setting, increased interferon signaling from mutations that stabilize the filament forming interfaces have potential to be pathogenic. Indeed, many *MDA5* gain-of-function mutations in MDA5 cause severe autoimmune disease (Rodero and Crow, 2016), although the mutations with clinical phenotypes reported so far map to the RNA and ATP binding sites. Hence the identification of the filament forming interface has predictive value, for researchers and also for clinicians who are likely to encounter patients with SNPs in the filament forming regions of MDA5.

### Structural changes in MDA5-dsRNA filaments coupled with ATP binding and hydrolysis

A comparative analysis of cryoEM structures determined in the presence of different ATP nucleotide analogs reveals clear correlations between the type and occupancy of the nucleotide bound in the catalytic site, the length of the RNA binding footprint and the helical twist of the filaments. Filaments formed with transition state analog ADP-AlF_4_ had a narrow distribution of intermediate twists (81° – 91°) and a 15-bp footprint, whereas filaments formed with a high concentration of ATP (10 mM) had mostly low helical twist (71° – 81°) and a 14-bp footprint (**Figure 1C**). In both of these cases the nucleotide occupancy in the active site was high (**Figure 3**). Filaments formed with lower concentrations of the nonhydrolyzable ATP analog AMPPNP (1 – 2.5 mM) had a broad range of twists (71° – 96°) and 15-bp footprints. Within these populations, the low-twist filaments (71° – 81°) had relatively high AMPPNP occupancy in the active site, but the filaments with intermediate and high twists had very little, if any nucleotide in the active site. The lack of AMPPNP density in these twist classes is not surprizing as the AMPPNP concentration present in the samples (1 – 2.5 mM) was of the same order of magnitude as the K_m_ for ATP reported for chicken MDA5 bound to a 24-bp dsRNA (2.2 ± 0.47 mM) (Uchikawa et al., 2016). Although the K_m_ (ATP) is likely to be smaller for MDA5 filaments on long dsRNAs, if we assume a K_d_ (AMPPNP) value of 1 mM, the occupancy of AMPPNP in the catalytic site can be expected to be 30 – 50% given the concentration of MDA5 in the imaged samples (3.45 µM). This would imply that less than half of the imaged MDA5 molecules contained AMPPNP in their active site. Consistent with this, more than half the particles formed with 1 – 2.5 mM AMPPNP had intermediate or high twist, lacked AMPPNP density in the active site, and had essentially identical structures as particles formed without nucleotide (**Figure 1**). We conclude that the ATP binding and hydrolysis are coupled to increases in the helical twist and RNA binding footprint of the filament.

Together, our structural data support the hypothesis that our cryoEM reconstructions have captured three distinct intermediates in the ATPase cycle. The low-twist structures correspond to the ATP-bound catalytic ground state, the intermediate-twist ADP-AlF_4_-bound structure is the transition state, and intermediate- and high-twist states that represent nucleotide-free states (**Figure 7D**). The coincidence of all three twist states on the same filament (**Figure 1B**) therefore suggests that multiple nucleotide binding states can coexist on one filament. The RecA-like domains Hel1 and Hel2 are in the closed conformation in the transition state and in the semi-closed conformation in the other states. The RNA binding footprint is 14 bp in the ground state and expands to 15 bp in the transition and nucleotide-free states. Notably, the increased twist and RNA footprint are maintained in the low nucleotide-occupancy states, even though the helicase domains return to the semi-closed state. Despite the increase in RNA footprint, the protein-RNA contact area decreases as the catalytic cycle proceeds, from 2,300 Å^2^ in the ground state to 2,100 Å^2^ in the intermediate-twist states and 1,900 Å^2^ in the high-twist nucleotide-free state. This implies that dissociation of ADP and Pi following hydrolysis causes MDA5 to loosen its grip on the RNA. Indeed, (Uchikawa et al., 2016) reached the same conclusion based on the crystal structures of MDA5 and LGP2 bound to dsRNA and different nucleotide analogs. Consistent with this, MDA5 forms longer, more continuous filaments in the presence of nonhydrolyzable ATP analogs than without nucleotide (Berke et al., 2012; Peisley et al., 2012; Wu et al., 2013). In contrast, binding of ATP or transition state analogs to RIG-I reduces the affinity of RIG-I for RNA (Rawling et al., 2015).

### Potential role of ATP hydrolysis in MDA5 function

The structural snapshots of the ATPase cycle we have obtained provide clues on the potential role of ATP binding and hydrolysis in MDA5 signaling, though open questions remain. The ATPase cycle of RIG-I and MDA5 has been proposed previously to perform a proofreading function in discrimination of self-versus non-self dsRNA by increasing the rate of dissociation of the protein from shorter endogenous RNAs (Lassig et al., 2015; Peisley et al., 2012; Rawling et al., 2015). In the case of RIG-I, ATP binding (Rawling et al., 2015) and hydrolysis (Lassig et al., 2015) have both been reported to promote dissociation from non-cognate RNA ligands. In the case of MDA5, ATP hydrolysis was found to enhance the binding specificity for long dsRNAs and promote formation of more continuous and stable filaments, while promoting dissociation from shorter dsRNAs (Peisley et al., 2012). As noted above, our structures show that binding of MDA5 induces significant distortions in the dsRNA backbone, and the extent and location of these distortions vary in the different intermediates of the ATPase cycle (**Movies S1-S5**). A possible interpretation is that ATP hydrolysis by MDA5 tests the physical properties of the RNA—specifically resistance to twisting and bending—such that MDA5 is more likely to remain associated with cognate ligands (exogenous long continuous RNA duplexes) and more likely to dissociate from non-cognate ligands (deaminated Alu repeats and short endogenous RNAs) due to the different way each type of ligand responds to the ATP-dependent conformational changes in MDA5, including changes in helical twist. This would provide a mechanical proofreading mechanism dependent on ATP hydrolysis. Consistent with this hypothesis, biochemical studies suggest that ATP binding contributes to proofreading by RIG-I by challenging the interaction with RNA and promoting dissociation (Rawling et al., 2015).

Our structural data indicate that ATP hydrolysis is coupled with a 1-bp expansion in the dsRNA binding footprint through the ratchet-like movements of Hel2 relative to Hel1 (**Figure 7B**). This could in principle result in translocation of dsRNA, however translocation would require cooperative binding of ATP to adjacent protomers and sequential hydrolysis in one direction along the filament. It appears more likely that expansion of the RNA binding footprint is a local phenomenon. Local expansion of the MDA5 footprint would provide a possible explanation for the reported repair of MDA5 filament discontinuities through ATP hydrolysis (Peisley et al., 2012). Expansion of the binding footprint of a long continuous filament could also explain how MDA5 can displace viral proteins from dsRNA in an ATP-dependent, CARD-independent manner (Yao et al., 2015). However, further work is required to determine more specifically how ATP hydrolysis propagates structural changes through MDA5-dsRNA filaments and how these changes may contribute to the proofreading and antiviral effector functions of MDA5.

## Acknowledgments

CryoEM data were collected at the MRC Laboratory of Molecular Biology, the UK national electron Bio-Imaging Centre (eBIC) at Diamond Light Source (DLS), and the University of Cambridge Cryo-EM Facility at the Department of Biochemistry. We acknowledge DLS for access and support of the cryoEM facilities at eBIC under proposal EM17434 funded by the Wellcome Trust, MRC and BBSRC. We thank scientists at MRC Laboratory of Molecular Biology (MRC-LMB) for support, particularly Kai Zhang for his assistance with cryoEM sample preparation; Shaoxia Chen, Giuseppe Cannone and Christos Savva for their assistance with data collection at MRC-LMB; and Yiquan Tang in William Schafer’s group for providing HEK293T cells. We are grateful to Yuriy Chaban and Dimitri Chirgadze for providing assistance in using the microscopes at eBIC and the University of Cambridge, respectively. We thank Shabih Shakeel for advice and for providing a script to rescale images collected at eBIC. We thank Sjors Scheres and Shaoda He (MRC-LMB), and Ed Egelman (University of Virginia) for helpful discussions regarding image reconstruction with helical symmetry. We are grateful for access to the MRC-LMB Scientific Computing and University of Cambridge High Performance Computing facilities. We acknowledge members of the Modis lab for insightful discussions. This work was supported by a Wellcome Trust Senior Research Fellowship to Y.M. (101908/Z/13/Z). K.Q. was supported by the European Research Council (ERC) under the European Union’s Horizon 2020 research and innovation programme ERC-CoG-648432 MEMBRANEFUSION awarded to John A. G. Briggs.

## Author contributions

Q.Y. and Y.M. conceived the study. Q.Y. performed the cell signalling and filament formation assays. Q.Y., K.Q. and Y.M. collected the electron microscopy data, analysed the data and prepared the figures. Q.Y. and K.Q. performed the image processing and reconstruction. Y.M. built and refined the atomic models. Q.Y. and Y.M. wrote the manuscript with discussion and input from K.Q.

